# Rhytidome- and cork-type barks of holm oak, cork oak and their hybrids highlight processes leading to cork formation

**DOI:** 10.1101/2023.03.31.535027

**Authors:** Iker Armendariz, Unai López de Heredia, Marçal Soler, Adrià Puigdemont, Maria Mercè Ruiz, Patricia Jové, Álvaro Soto, Olga Serra, Mercè Figueras

## Abstract

The periderm is basic for land plants due to its protective role during radial growth, which is achieved by the polymers deposited in the cell walls. In most trees, like holm oak, the periderm is frequently replaced by subsequent internal periderms yielding a heterogeneous outer bark made of a mixture of periderms and phloem tissues, known as rhytidome. Exceptionally, cork oak forms a persistent or long-lived periderm which results in a homogeneous outer bark of thick phellem cell layers known as cork. Here we use the outer bark of cork oak, holm oak, and their natural hybrids’ to analyse the chemical composition, the anatomy and the transcriptome, and further understand the mechanisms underlying periderm development. The inclusion of hybrid samples showing rhytidome-type and cork-type barks is valuable to approach to cork and rhytidome development, allowing an accurate identification of candidate genes and processes. The present study underscores that biotic stress and cell death signalling are enhanced in rhytidome-type barks whereas lipid metabolism and cell cycle are enriched in cork-type barks. Development-related DEGs, showing the highest expression, highlight cell division, cell expansion, and cell differentiation as key processes leading to cork or rhytidome-type barks.

## INTRODUCTION

The periderm arises during radial thickening of stems and roots (secondary growth) and confers protection against water loss and pathogen entrance and overall contributes to the plant fitness (Serra *et al*., 2022). This protective function is afforded by the phellem, which accumulates a lignin-like polymer and a suberin polyester in their cell walls. The periderm is important in herbaceous, tubers and some fruits but specially in woody plants, where it is the prevalent protective tissue. In woody species, new phloem (bast) is produced outwardly and new xylem inwardly from the vascular cambium every year during the growing season (Tonn and Greb, 2017). This newest xylem and phloem push the outer layers centrifugally and a new phellogen is formed within the area of the older phloem, protecting the young phloem from outside (Howard, 1977). Like vascular cambium, phellogen or cork cambium is a bifacial and lateral meristem activated seasonally. Periclinal divisions of phellogen cells produce phellem outwardly and phelloderm inwardly. The structure formed by phellem (cork), phellogen and phelloderm constitute the periderm (Evert *et al*., 2006). In most woody species, and contrarily to vascular cambium, phellogen has limited activity, and successive phellogens differentiate in inner positions in the bark. When a new periderm is formed inward, the outer tissues including the older periderm will eventually die (Howard, 1977). The newest phellogen marks the limit of the inner bark (comprising the living phloem) and the outer bark, the later usually forming a so-called rhytidome (Romberger *et al*., 1993). This rhytidome therefore includes successive thin, suberized and intricate phellem layers, enclosing heterogeneous cortical tissues (parenchyma, fibres, etc.) and collapsed phloem cells (De Burgos *et al*., 2022).

Noteworthy, the phellogen is thought to be active throughout the tree life in cork oak (*Quercus suber*) (Silva *et al*., 2005), and as such, it forms a persistent or long-lived periderm (Serra *et al*., 2022). Therefore, there is a unique, thick, and continuous periderm mostly consisting of phellem cells known as cork. Cork has economic and environmental relevance. It is an industrially profitable renewable raw material and suberin recalcitrance elicits CO_2_ sequestration, which is favoured by the periodic extraction of cork that stimulates the cork production between 250 and 400% (Gil, 2014). Despite the uniqueness of cork oak in maintaining a persistent periderm, the cellular and molecular mechanisms that trigger its persistence by encompassing the internal growth are still largely unknown. Previous transcriptomic studies of outer barks of cork oak and rhytidome-developing oaks (*Q. ilex* and *Q. cerris*) highlighted some processes and genes enriched in rhytidome and cork but the identification of differentially expressed genes was limited due to the low-coverage offered by Roche-454 Life Sciences platform and by the lack of biological replicates (Boher *et al*., 2018; Meireles *et al*., 2018).

Cork oak shares habitat and hybridises naturally with holm oak (*Quercus ilex*) (Burgarella *et al*., 2009), a species showing the typical rhytidome. *Q. ilex* x *Q. suber* offspring differ in their outer bark anatomy, although generally they show a rhytidome-like outer bark, similar to *Q. ilex* but with significantly thicker phellem layers (De Burgos *et al*., 2022). Our aim in this study is to identify the molecular mechanisms underlying the formation of the two main bark types, rhytidome and cork as a “single thick phellem”. For this purpose, we have included in our transcriptomic analysis not only *Q. ilex* (rhytidome) and *Q. suber* (cork) samples, but also hybrid individuals, with different introgression levels and intermediate barks. Using the Illumina platform and the availability of cork oak genome, the comparison of cork-type and rhytidome-type barks transcriptomes provides new candidate genes of cork formation related to development, cell division, growth and differentiation.

## MATERIALS AND METHODS

### Outer bark harvesting

We harvested outer barks of tree trunks from four adult cork oaks (*Quercus suber* L.), four holm oaks (*Quercus ilex* L.) and six *Q. ilex* x *Q. suber* hybrids. Trees were naturally grown in a mixed holm oak-cork oak forest in Fregenal de la Sierra (Extremadura, Spain). These hybrids were previously identified according to morphological features and molecular markers and a detailed anatomy was reported recently (López de Heredia *et al*., 2020; De Burgos *et al*., 2022). The samples were obtained when the phellogen activity was high enough to allow the outer bark detachment from the inner bark. Outer bark was harvested from the south-facing part of the trees, at breast height, and it was manually removed from the trunk using a hammer and a chisel. The material was collected from the inner face of the outer barks scratching with a chisel, immediately frozen in liquid nitrogen and kept at -80 ºC for further use. Anatomical observations were performed as detailed in De Burgos *et al*., (2022).

### Chemical analysis of the outer barks

Chemical analyses were performed in one representative sample of cork and rhytidome, five samples of rhytidome-like bark hybrids and one sample from the cork-like bark hybrid (FS1) identified. The summative chemical analyses included the determination of ash, extractives, suberin, Klason lignin and holocellulose. The ash content was determined by incinerating 2 g of cork at 525ºC during 1 h with a muffle furnace (Faenza, Italy). Extractives were determined by successive Soxhlet extraction with dichloromethane (6 h), ethanol (8 h) and hot water (20 h).

After each extraction, the cork residue was air-dried and kept for subsequent analysis and the extracted solution was evaporated to obtain the solid residue, which was weighed. The suberin content was determined in extractive free material by alkaline methanolysis for its depolymerisation using a Soxhlet in reflux mode during 3 h. Then, the extracted liquid was acidified with 2 M H_2_SO_4_ to pH 6, and evaporated to dryness in a rotating evaporator (Aircontrol, Spain). This residue was suspended in 100 ml H_2_0 and extracted with 100 ml CHCl_3_ three times. The combined extracts were dried over Na_2_SO_4_, filtered, evaporated, and determined gravimetrically as suberin. On the other hand, the desuberized solid material was used for Klason lignin determination by a hydrolysis with 72% H_2_SO_4_ (Jové *et al*., 2011). Holocellulose fraction was isolated from desuberized fraction by delignification using acid chloride method (Wise et al. 1946). All measurements were reported as a percentage of the original sample. Principal components analysis (PCA) was performed to plot the variation of outer bark chemical composition using the log-transformed data of the percentage of each fraction (variables) in the eight samples.

### Total RNA extraction and purification

Total RNA was extracted from outer barks using a modified method described previously (Chang *et al*., 1993; Chaves *et al*., 2014). Two grams of tissue were grounded in liquid nitrogen using a mortar and pestle and rapidly mixed with 15 ml of preheated (65 ºC) CTAB extraction buffer (2% CTAB, 4% PVP-40, 300 mM Tris-HCl pH 8.0, 25 mM EDTA, 2 M NaCl, and 3.3% 2-mercaptoethanol) using a vortex. After a 10 min incubation at 65 °C, we extracted the sample twice with one volume of chloroform: isoamyl alcohol 24:1 (v:v) followed by centrifugation each at 15,000 *g* for 20 minutes. The aqueous fraction was precipitated using 1 V of isopropanol and 0.1 V of NaOAc 3 M (pH 5.2) and incubated for 3 h at -80 °C or overnight at -20 °C. The precipitate was collected by centrifugation at 15,000 *g* for 30 min, resuspended in 700 μl of preheated (65 °C) SSTE buffer (1 M NaCl, 0.5% SDS, 10 mM Tris-HCl pH 8 and 1mM EDTA) and treated twice with the same volume of chloroform: isoamyl alcohol (24:1 (v:v)) and centrifuged 10 min at 21,000 *g*. The supernatant was precipitated overnight with 2 V of ethanol 100% at -80 °C and collected by centrifugation at 21,000 g for 30 min at 4 ºC. After two washes of 70% ethanol, the nucleic acid pellet was resuspended in 50 μl of RNase-free water. RNeasy Power Plant Kit (Qiagen) and DNAse I on-column digestion were used to remove polyphenols and genomic DNA, respectively. As we had already the extracted RNA, we adapted the procedure of the commercial kit by adding 500 μl of MBL (99: 1 MBL: β-mercaptoethanol) to 50 μl of each sample together with 50 μl of PSS and 200 μl of IRS. The total RNA yield was measured with a Nanodrop and the RNA integrity values (RIN) were obtained with a Bioanalyzer 2100 (Pico RNA 6000 Kit, Agilent). The values obtained for each of the samples are shown in **Supplementary Table S1**.

### Analysis of RNA-seq high-throughput mRNA sequencing data

Outer bark cDNA libraries were obtained using the MGIEasy RNA Library Prep Kit V3.1 and 3 μg of each sample (RIN value > 8). Sequencing was performed by the BGISEQ500 (paired-end reads of 100 bp) at BGI Genomics (Hong Kong). In total, 16 samples were sequenced. For each cork oak, holm oak and rhytidome-like groups, four south-oriented bark samples each from a different tree were sequenced. For the cork-like hybrid unique individual, 4 bark samples extracted from the north, south, west, and east orientations were sequenced. A minimum of 100 M reads was obtained for each library. The quality of raw reads was assessed with FASTQC software (Andrews, 2010) and removal of the first low-quality 12 bp was performed with Trimmomatic (Bolger *et al*., 2014). The reads were mapped with GSNAP (Wu et al., 2016) against *Q. suber* genome as a reference (GCF_002906115.1_CorkOak1.0_genomic.fna) (Ramos *et al*., 2018), and the unique concordantly mapped reads were kept for library construction. Reads from each library were assembled with Cufflinks and the consensus transcriptome for all the samples was generated assembling each library with Cuffmerge (Trapnell *et al*., 2010, 2012). Willing to work with unique gene identifiers, different isoforms were collapsed using the genome positions and total counts were estimated by HTSeq-count (Anders *et al*., 2015). PCA analysis of transcript profiling was conducted and represented using DESeq2 (Love *et al*., 2014), by considering the variation of rlog data from 47,292 gene loci in the 13 outer bark samples extracted from south orientation. The results are displayed in bivariate diagrams showing the main factors displayed by ggplot2 (Wickham, 2016).

The count matrix was generated using the 16 libraries and allowed to identify differentially expressed genes (DEGs) using the DESeq2 package (Love *et al*., 2014). Read counts per locus were corrected by rlog transformation and DEGs were obtained by pairwise comparisons: cork (cork oak bark) *vs* rhytidome (holm oak bark), cork *vs* cork-like bark (hybrid bark similar to cork), cork *vs* rhytidome-like bark (hybrid bark similar to rhytidome), cork-like *vs* rhytidome, rhytidome-like bark *vs* rhytidome and cork-like *vs* rhytidome-like. Transcripts with an adjusted p-value smaller than 0.01 and log2FC ≤ -1 and ≥ 1 were considered as DEGs. The normalized count data were used for hierarchical clustering of DEGs using the MeV program (Howe *et al*., 2011) by k-means and Euclidean distance with 100,000 iterations. For *Arabidopsis thaliana* annotation we used the Blastp and the TAIR10 library from Ensembl, with the options num_alignments 1 and evalue 1e^-08^. AgriGO V2.0 (Tian *et al*., 2017) was used for gene ontology enrichment for the best Arabidopsis homologs (FDR ≤ 0.05). The GO terms were manually collapsed based on the analogous description and the set of genes they contained.

### Real-time quantitative PCR

The analysis was performed in all biological replicates using the primers of six genes (**Supplementary Table S2)**. First-strand cDNA was synthesized from 200 ng DNase digested RNA using RevertAid First Strand cDNA Synthesis Kit (Thermofisher). The synthesis of cDNA was performed using oligodT primer and following manufacturer’s instructions. The program for the cDNA synthesis was as follows: 16°C for 30 min; 60 cycles of 30°C for 30 s, 42°C for 30 s and 50°C for 60 s; 85°C for 5 min. Real-time PCR analysis was performed using a LightCycler® 96 Real-Time PCR System (Roche). Primers were designed for each gene with Primer3-0.4.0 software (http://bioinfo.ut.ee/primer3-0.4.0/). Each RT-qPCR reaction (10 μl) contained 5 μl of SYBR Green Select Master Mix (Roche), 300 nM of the corresponding forward and reverse primers, and 2.5 μl of a 25-fold diluted cDNA. The conditions of the thermal cycle were the following: 95 °C for 10 min; 40 cycles of 95 °C for 10 s and 60 °C for 60 s. A final dissociation step of 85 °C for 5 min was included to confirm a single amplicon. For each primer pair, standard curves with a five-fold dilutions series of a cDNA mix corresponding to equal amounts of all biological replicates of cork bark, rhytidome bark, cork-like bark, and rhytidome-like bark (1/10, 1/25, 1/50, 1/100, and 1/250) were used to determine amplification efficiency of each gene (E = 10 ^(−1/slope)^). The mRNA abundances for each gene were calculated as relative transcript abundance = (E_target_)^ΔCt target (control-sample)^ / (E_reference_)^ΔCt reference (control-sample)^ (Pfaffl, 2001). The calibrator or control sample consisted of equal amounts of cDNA of all biological replicates. The housekeeping gene used to normalize the results was tubulin (Soler *et al*., 2008). DNA contamination was ruled out at the beginning and controls to confirm no presence of environmental contamination were included in each experiment. Three technical replicates were used for every four biological replicates.

## RESULTS

### Anatomical and chemical analyses to classify the outer barks of *Q. suber* x *Q. ilex* hybrids and the parental lines

Microscopic observations of cross-sections under UV light after phloroglucinol staining highlighted the suberized cell walls of the different outer barks used in this study (**Fig. 1**). The outer bark of *Q. ilex* represented the typical rhytidome displaying thin periderms consisting of few phellem cell layers (**Fig. 1A**). In contrast, *Q. suber* outer bark showed a single periderm consisting of a thick and homogeneous tissue based on suberized phellem cells (**Fig. 1B**). Most of the natural hybrids identified previously were categorized as F1 hybrids through genetic analysis (López de Heredia *et al*., 2020). They showed a rhytidome, similar to that of *Q. ilex*, but with closer and thicker periderms and additionally, some of them presented a singular suberization of inactive phloem between periderms (**Fig. 1C**). On its side, the FS1 hybrid had a unique outer bark phenotype with much thicker phellems, rather like those of cork oak (**Fig. 1D**). In agreement, this FS1 hybrid was identified previously as a backcross with *Q. suber*. Due to this phenotype similarity, this cork-producing hybrid (FS1 hybrid) will be referred to as a cork-like hybrid and the remaining hybrids as rhytidome-like hybrids.

**Fig. 1.**
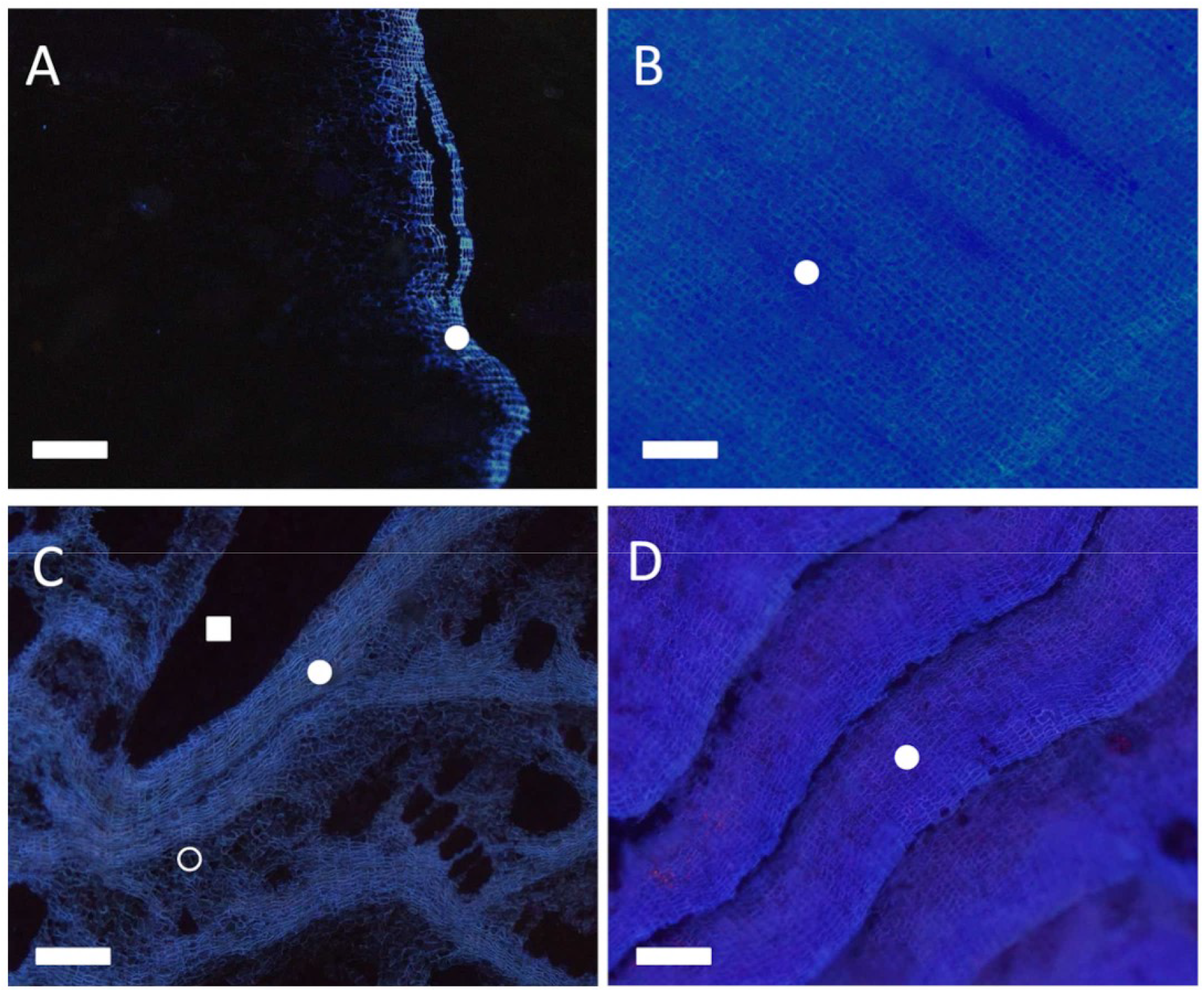
Outer bark anatomy of cork oak, holm oak and their hybrids. Suberized cell wall fluorescence detected in cross-sections under UV light after phloroglucinol-HCl staining. A) Holm oak (*Q. ilex*), B) cork oak (*Q. suber*), C) F1 hybrid with rhytidome-like phenotype, D) FS1 specific hybrid backcrossed with *Q. suber* and with a cork-type phenotype. Phellem layers (closed circle), suberized inactive phloem (open circle) and a lignified phloematic ray (closed square). Scale bars: 200 μm.

Consistently with these observations, the chemical analyses of the outer barks make it possible to distinguish different groups based on the proportion of the different components (holocellulose, lignin, suberin and water-, ethanol- and dichlorometane-soluble extractives). The cork and rhytidome-type (rhytidome-like and rhytidome) were at opposite ends of the first principal component axis of the PCA, which explained 69% of the variance (**Fig. 2A**). In this axis, cork-like was found between cork and rhytidome-type. A more detailed inspection of the data showed a gradient in the percentage of suberin and dichlorometane extractives in the bark samples. The cork and cork-like FS1 outer bark samples had a ten-fold higher percentage of suberin than barks of holm oak and rhytidome-like hybrids (**Supplementary Table S3, Fig. 2B**), in agreement with cork oak and holm oak outer bark composition reported previously (Holloway 1983). Concomitantly, cork and cork-like FS1 samples presented a boost in the proportion of dichloromethane extractives, which contained non-polar components such as terpenes and waxes. The abundance of both types of compounds agrees with the common fatty acyl precursors of suberin and waxes (Li *et al*., 2007; Serra *et al*., 2009*a*). Conversely, the outer bark of holm oak and the rhytidome-like hybrids contained on average 2.7 times more ethanol-soluble extractives than cork and cork-like outer barks. Interestingly, the holocellulose percentage was 3-fold higher in holm oak and all the hybrids (including the cork-like sample) than in the cork oak bark.

**Fig. 2.**
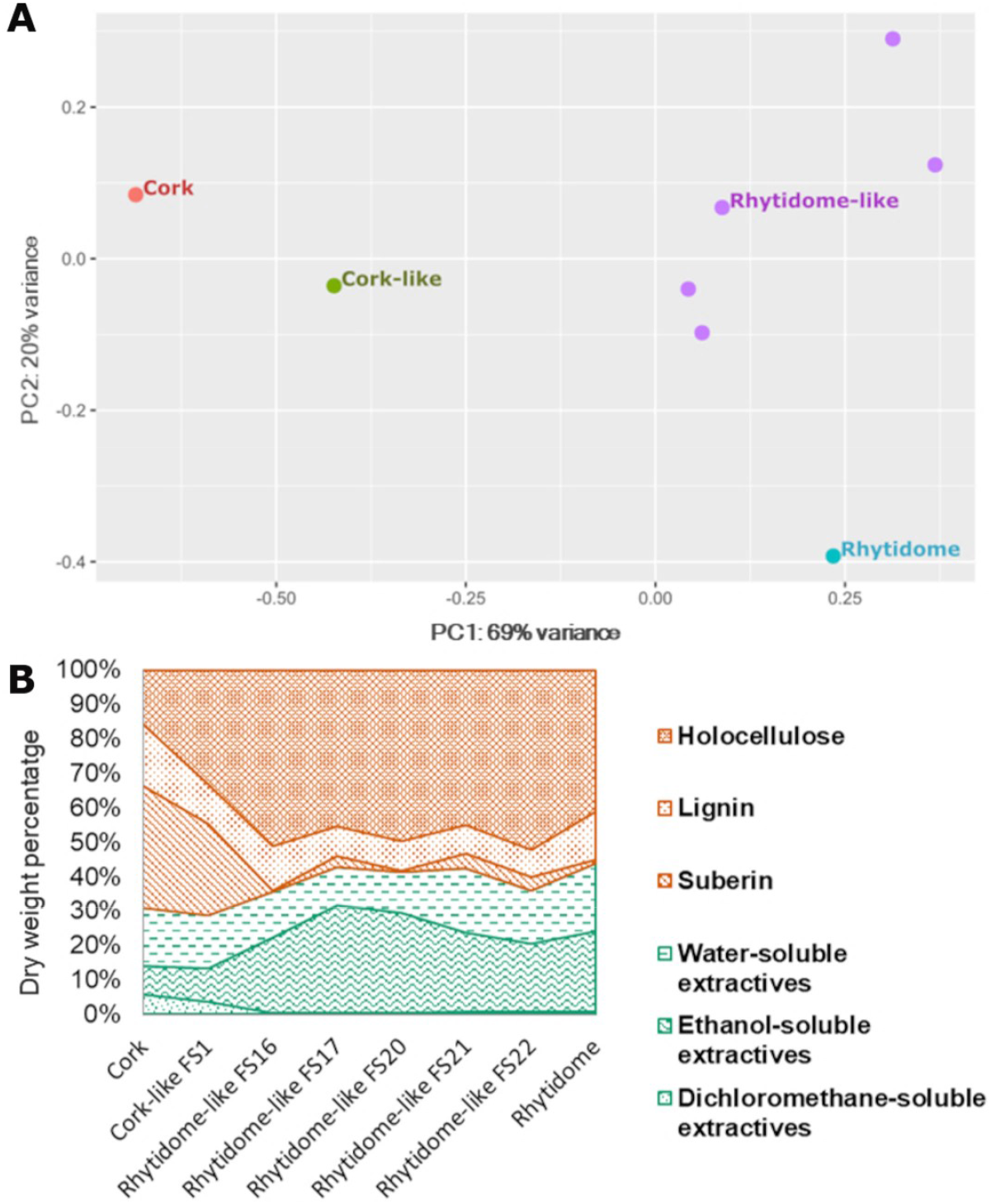
Chemical composition of the outer barks of cork oak, holm oak and their hybrids. A) Principal component analysis (PCA) of the data from chemical composition analysis of the outer barks of cork oak, holm oak and the hybrids. The first principal component shows a clear separation between cork-type and rhytidome-type barks and a gradient between cork, cork-like hybrid and the rhytidome-type barks. B) Dry weight percentage of the outer bark chemical composition of cork oak, holm oak, and a set of hybrids showing rhytidome-like bark and the hybrid showing a cork-like bark. Note the higher relative percentage of suberin and dichloromethane-soluble extractives in the cork-type barks.

### Cork- and rhytidome-type barks have the most different transcriptomes

To understand the molecular processes that differentiate the outer bark producing cork from others that generate the typical oak rhytidome, we sequenced the transcriptomes of the outer bark from *Q. suber, Q. ilex*, and their natural hybrids (*Q. ilex* x *Q. suber*). The raw reads obtained for each library were pre-filtered to remove adaptors, contaminants, and low-quality reads. Statistical results of processed data are shown in **Supplementary Table S1**. On average, 82.61% of the reads mapped uniquely and concordantly against the cork oak genome (GCF_002906115.1_CorkOak1.0) (Ramos *et al*., 2018) and consensus transcriptome covered 47,292 different transcripts, corresponding to 16,192 Arabidopsis (TAIR10) protein matches. The distance analysis of global transcript profile showed the highest similarity between biological replicates and also high similarity of cork-like outer bark with cork replicates (**Supplementary Fig. S1**). PCA analysis of the transcriptomes showed that the first principal component explained 57% of the total variance and distributed the outer bark types in a gradient from cork to rhytidome, with parental lines at each side (**Fig. 3A**).

**Fig. 3.**
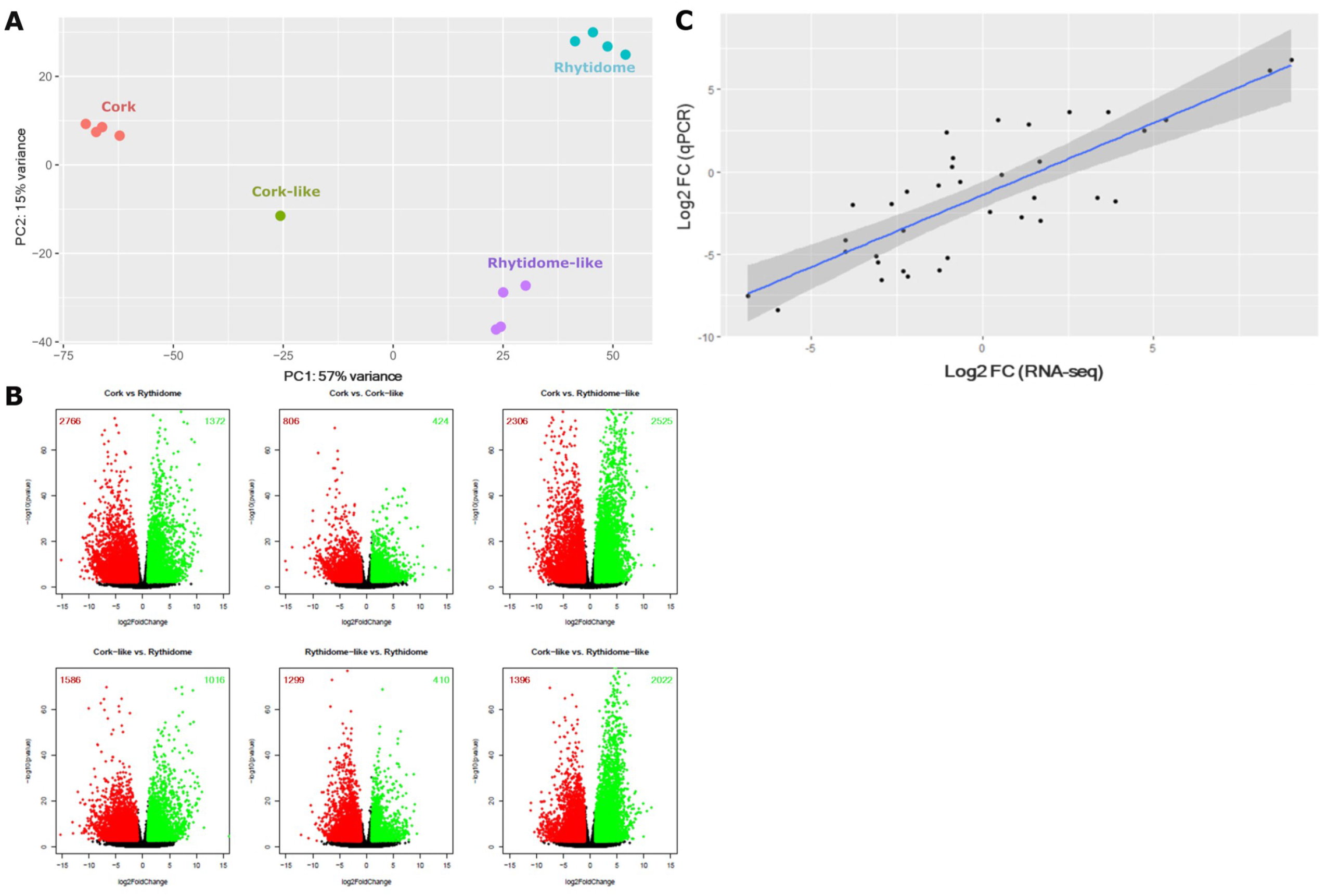
Transcriptome profile and differential expression analysis of the different outer barks. A) Principal component analysis of the global transcript profile obtained from the outer barks of cork oak, holm oak and the hybrids. Similar transcriptomes within individuals of the same bark-type group together. The first principal component shows a clear separation between cork-type and rhytidome-type barks, as well as a gradient between cork, hybrids and rhytidome outer barks. B) Volcano plot showing odds of differential expression (-log10 p-adjusted value) against ratio (log2 FoldChange) of different pairwise comparisons: cork/rhytidome, cork/cork-like, cork/rhytidome-like, cork-like/rhytidome, rhytidome-like/rhytidome, cork-like/rhytidome-like. Genes with –log10 greater than 2 and with log2FC absolute value greater than 1 are considered as DEGs. Green dots depict upregulated genes and red dots downregulated genes for each comparative. The number of upregulated and downregulated genes found in each comparison are shown in green and red, respectively within each graph. C) Correlation graph of the mRNAs log2ratio values between the RNA-seq and the qPCR analyses. The Pearson correlation coefficient (ρ) is 0.804 and the p-value < 0.001 (3.43 10^−9^). The shaded area represents the confidence interval of the regression line.

We next identified the differentially expressed genes (DEGs) performing a pairwise comparison between transcriptomes of each bark types. We considered DEGs those genes with a padj < 0.01 and a log2FC either < -1 or > 1 (**Fig. 3B**). Overall, we found 8,336 DEGs at least in one of the comparisons (**Supplementary Table S1; Fig. 3B**). Volcano plots showed that the comparisons with the largest number of DEGs, and thus more divergent samples, are cork/rhytidome-like (4,831 DEGs) and cork/rhytidome (4,138 DEGs) (**Fig. 3B**). Conversely, the comparisons presenting the lowest number of DEGs were cork/cork-like (1,230 DEGs) and rhytidome-like/rhytidome (1,709 DEGs). This is consistent with the anatomical and chemical phenotypic similarity of hybrids with their corresponding parental lines.

To validate the RNA-seq data, six genes (**Supplementary Table S2**) were analysed in the five comparisons by RT-PCR. Log2ratio of RTA values were compared to Log2FC of RNA-seq data (**Fig. 3C**). The statistical analysis presented a Pearson correlation coefficient of 0.804 and a p-value < 0.001 (3.43 10^−9^), hence indicating a positive correlation between PCR and RNA-seq results.

To identify the functional networks of proteins that distinguish bark types, we clustered the co-regulated genes and predicted the enriched functional processes for each cluster. Based on their expression pattern, DEGs grouped in eight clusters (**Fig. 4**) that we classified into: (i) clusters with gene expression biased toward rhytidome-type barks (cluster 1, cluster 2 and cluster 3), (ii) clusters with gene expression biased toward cork-type barks (cluster 4, cluster 5, cluster 6 and cluster 8) and (iii) a cluster (cluster 7) with maximum and opposite expression in cork-type samples – downregulated in cork and upregulated in cork-like bark samples.

**Fig. 4.**
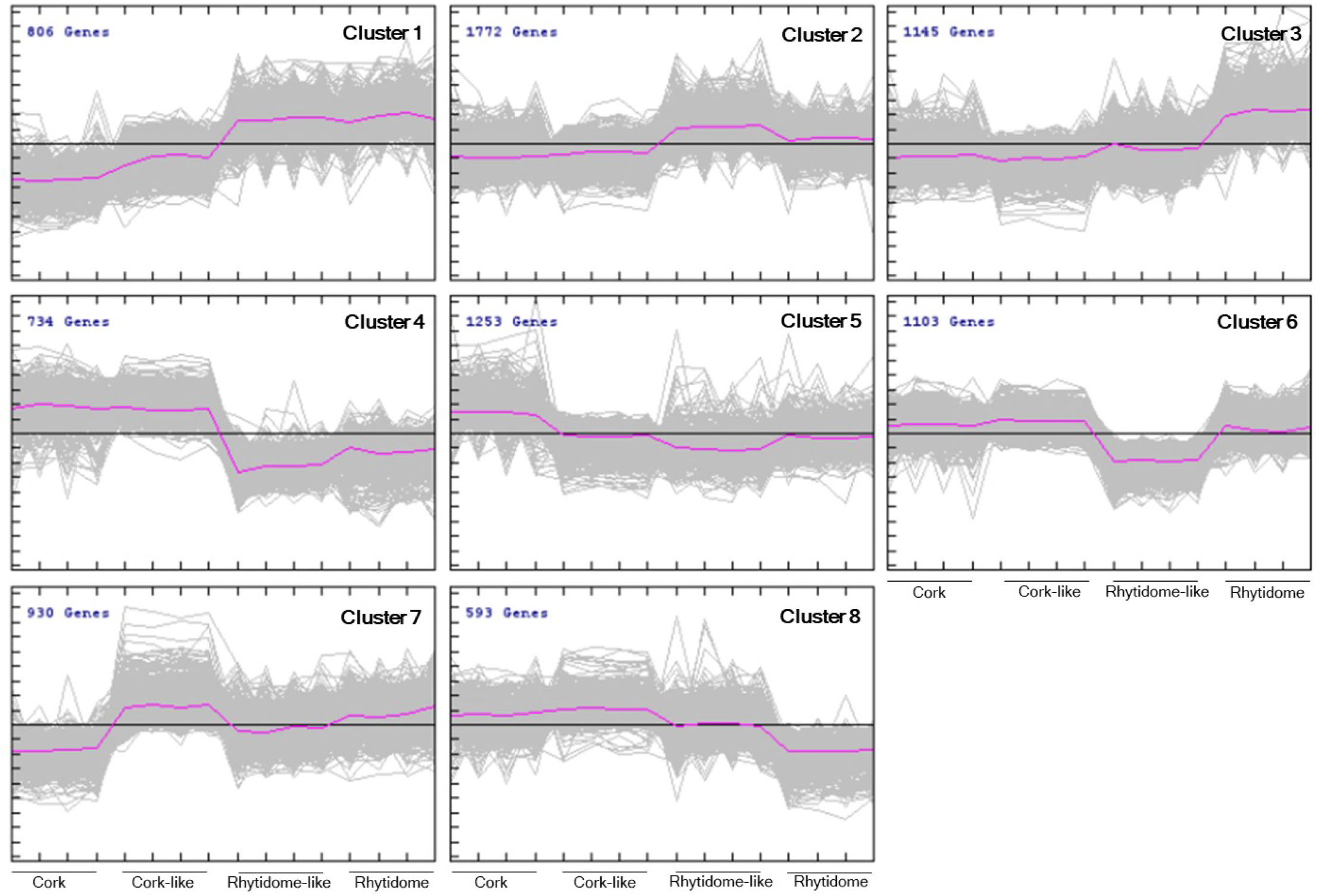
Cluster analysis of DEGs according to their expression profile in the different outer bark types. Eight clusters were obtained. Each cluster panel shows the number of genes included and the individual and averaged gene expression profile (rlog), in grey and purple lines, respectively. also shown. Clusters 1, 2, 3 contain genes upregulated in rhytidome-type outer barks. Clusters 4, 5, 6, and 8 contain genes upregulated in cork-type barks. Cluster 7 shows particular expression peaks in cork-like and rhytidome outer barks.

### Rhytidome-type barks are enriched in abiotic and biotic stress, phenylpropanoid metabolism, development and cell death

Clusters with gene expression biased toward rhytidome-type barks were cluster 1, cluster 2 and cluster 3. Cluster 1 included 9.67% of DEGs with highest expression in rhytidome and rhytidome-like barks. It was enriched in gene ontologies related to the following biological process: (i) abiotic and biotic stress-related signalling, (ii) regulation of transcription, (iii) phenylpropanoid metabolism: lignin and flavonoid biosynthesis, (iv) developmental process, (v) hormone metabolism, (vi) pigment biosynthesis and (vi) cell death (**Supplementary Fig. S2, Supplementary Table S4**). Cluster 2 included the largest number of DEGs (21.26%) that showed a peak of expression in rhytidome-like bark. In this cluster, we found enrichment in the GO terms associated with (i) response to abiotic stress, (ii) RNA metabolism and gene expression, (iii) aromatic compound metabolism, (iv) development and reproductive development, (v) energetic metabolism, (vi) cell communication and signalling, (vii) circadian rhythm and (viii) photoperiod and flowering (**Supplementary Fig. S2, Supplementary Table S4**). Cluster 3 held 13.74% of DEGs, which were upregulated in the rhytidome bark. The enriched biological processes of this cluster were: (i) response to biotic and abiotic stimuli, (ii) response to hormones, (iii) immune system, (iv) protein phosphorylation, (v) transport (related with organic and inorganic compounds and membrane and non-membrane dependent transport), (vi) biogenesis of the cell wall, (vii) processes of secondary metabolism, mostly related with phenylpropanoids and lignin, (viii) cell death, (ix) development, (x) biosynthesis of jasmonic acid and (xi) transcription regulation (**Supplementary Fig. S2, Supplementary Table S4**). Hence, rhytidome-type bark induces expression of genes related to abiotic and biotic stresses, phenylpropanoid metabolism, development and cell death.

### Genes upregulated in cork-type barks are related to lipid and phenylpropanoid metabolism, development and cell wall biogenesis

Clusters with gene expression biased toward cork-type barks were clusters 4, 5, 6 and 8. Cluster 4 contained 8.81% of DEGs and represented genes upregulated in both cork-type barks. The biological processes enriched in this cluster were: (i) lipid metabolism, (ii) organic acid metabolism, (iii) oxidation-reduction processes, (iv) secondary metabolism, mostly related with lignin and suberin biosynthesis, (v) carbohydrate metabolism, (vi) processes related with cell wall biogenesis and organization, (vii) coenzyme metabolism, (viii) steroid metabolism, (ix) cutin biosynthesis, (x) response processes and (xi) cuticle development (**Supplementary Fig. S3, Supplementary Table S4**). Cluster 5 represented the 15.03% of DEGs which were induced in cork bark and repressed in rhytidome-like bark and presented the greatest diversity in GO terms. Processes enriched in this cluster were: (i) primary metabolism, (ii) development and cell cycle, (iii) carbohydrate metabolism, (iv) carboxylic acid metabolism, (v) lipid metabolism, (vi) response to stimuli, (vii) DNA metabolism, (viii) cell morphogenesis and cell wall organization, (ix) microtubule-dependent processes, (x) cofactor metabolism, (xi) nitrogen bases metabolism and phosphorylation, (xii) phosphorous-containing compound metabolism, (xiii) oxidation-reduction processes, (xiv) hydroxyl compound metabolism, (xv) phenylpropanoid metabolism, (xvi) steroid metabolism and (xvii) vesicle-mediated transport (**Supplementary Fig. S3, Supplementary Table S4**). Cluster 6 contained 13.23% of DEGs and these were upregulated in cork-type bark and downregulated in rhytidome-like bark. This cluster displayed enrichment in GO terms related to (i) response to abiotic stress, (ii) secondary metabolism, mostly related with lignin, (iii) protein phosphorylation, (iv) cell wall organization and biogenesis, (v) carbohydrate metabolism, (vi) cell development, (vii) cell cycle, (viii) cell communication and (ix) oxidation-reduction processes (**Supplementary Fig. S3, Supplementary Table S4**). Cluster 8 included the lowest number of DEGs (7.11%) and showed upregulation in cork-type barks and downregulation in rhytidome bark. In this cluster, the only biological process enriched was the response to stimulus (**Supplementary Fig. S3, Supplementary Table S4**). In conclusion, these clusters are committed to processes related to lipid and phenylpropanoid metabolisms, development and cell wall biogenesis.

### Cluster 7 is a cluster strongly upregulated in cork-like bark and downregulated in cork bark

Cluster 7 encompassed 11.16% of DEGs, which were upregulated in cork-like and rhytidome barks and strongly downregulated in cork bark (**Supplementary Fig. S4, Supplementary Table S4**). Within this cluster, the biological processes enriched were: (i) response to biotic and abiotic stimuli, (ii) photosynthesis, (iii) protein phosphorylation, (iv) carboxylic acid metabolism, (v) cell death, (vi) senescence, and (vii) pollen recognition. Consistently with oxidative stress associated with biotic and abiotic stress and photosynthesis, there is also a group of genes related to reactive oxygen species metabolism. The enrichment of cell death and senescence in this cluster points out that genes found in these two processes are enhanced in cork-like and rhytidome barks in comparison to cork bark.

### Stress, development, secondary metabolism and cell wall metabolism can be found in cork and rhytidome-type barks

Inspecting over the processes enriched in clusters classified as rhytidome-type or cork-type (above sections), we observed processes that were commonly or specifically enriched, which may bring interesting information. For example, most clusters (1, 2, 3, 5, 6, 7 and 8) highlighted abiotic stress response, hence supporting that phellem formation is related to abiotic stress signalling. Biotic stress and defence were found enriched in clusters 1, 2, 3, 4 and 7 as well. Secondary metabolism and specifically phenylpropanoid metabolism were found in clusters with opposite behaviour (1, 3, 4, 5, and 6). Despite lipid metabolism was enriched in clusters 4 and 5 (high expression in cork-type), we identified genes related to suberin biosynthesis or its upstream pathways (Aralip database: fatty acid synthesis, fatty acid elongation, and wax biosynthesis, **Supplementary Table S1**) in all clusters, but cluster 4 had the highest ratio of suberin-related genes (7.35% of total genes of the cluster), followed by clusters 5 and 6 (1.83% and 1.81% respectively). These results are consistent with the anatomy and chemical composition as suberin synthesis takes places in both rhytidome and cork-type barks. Development and cell wall biogenesis were also GO terms found in clusters with opposite bark-type (Development: clusters 1, 2, 3, 4, 5 and 6, and cell wall biogenesis: clusters 3, 4, 5, and 6) suggesting that these genes could account for the differences involved in bark development. Finally, cell death was enriched in clusters 1, 3, and 7, which had in common gene downregulation in cork bark and therefore suggested that the lack cell death could contribute to the cork bark features.

## DISCUSSION

Analysis of the chemical composition of the outer bark regarding holocellulose, suberin, lignin and extractives content yielded results consistent with anatomical observations. The transcriptome comparison using outer barks showing cork or rhytidome features provided 8,336 DEGs, including those identified in hybrid individuals. Genes clearly upregulated in rhytidome-type barks were found in clusters 1, 2 and 3, while genes upregulated in cork-type barks were in clusters 4, 5, 6, and 8. Clusters 2 and 7 were specifically upregulated in hybrid individuals, with DEGs upregulated in rhytidome-like bark hybrids (cluster 2) or in cork-like bark hybrid (cluster 7).

Hybridization and introgression are well known to modify gene expression, due to the disruption of regulation pathways, mainly of trans-acting regulators, epistatic relationships or the lack of intermediate gene products acting in complex metabolic routes, for example (Jin *et al*., 2008; Czypionka *et al*., 2012; Liang *et al*., 2018; Silvert *et al*., 2019; Kong *et al*., 2020). This is the case of bark development, where F1 hybrids, carrying a copy of *Q. suber* genes, fail to form a long-living or persistent periderm. Maybe more interesting is the general suberization of inactive phloem, suggesting an alteration of expression patterns in this tissue, prior to its final death (de Burgos et al., 2022). Genes upregulated specifically in rhytidome-type hybrids (cluster 2), corresponding to GOs related to response to abiotic stress, RNA metabolism and gene expression, aromatic compound metabolism and development, could underlie this feature.

On the other side, the individual identified as a backcross with cork oak, the cork-like bark hybrid FS1 (López de Heredia et al., 2020), is expected to carry, on average, two alleles coming from cork oak on half of the genes involved in bark formation. Consistently, it showed much thicker layers of phellem in its outer bark, while no suberization of inactive phloem had been detected. The clues of this thicker phellem could be in cluster 4 and 6, which corresponded to GOs related to lignin, suberin, cell wall formation, cell development and cell cycle. Cluster 7 showed the most differential features between cork and cork-like bark hybrid, which GOs were biotic and abiotic response and signalling, cell death and senescence among others.

### Outer bark development: the most highly expressed genes give some clues about the differential features between cork and rhytidome-type barks

Among the most expressed genes related to developmental process in rhytidome-type bark clusters (**Supplementary Table S5**) stood out genes related to periderm development in Arabidopsis root (ARF6), suberin monomers transport (ABCG11), epidermal cell morphology (Myb5), protophloem and xylem cell differentiation (Bam3 and KNAT1, respectively), cell expansion reduction (Feronia), flowering delay (Frigida-like genes), repression of cell division during flower organ growth (ARF2), programmed cell death (RRTF1/ERF109) and organ abscission (SOBIR1) (Michaels *et al*., 2004; Schruff *et al*., 2006; Li *et al*., 2009; Panikashvili *et al*., 2010; Depuydt *et al*., 2013; Haruta *et al*., 2014; Liebsch *et al*., 2014; Bahieldin *et al*., 2016; Taylor *et al*., 2019; Xiao *et al*., 2020). Moreover, we found other genes highlighted as relevant for vascular patterning such as STM, SVP, PTL, LBD4, and LBD1 (Yordanov *et al*., 2010; Liebsch *et al*., 2014; Zhang *et al*., 2019; Smit *et al*., 2020) and relevant to meristem activity and even cambium activity such as CLV1, CLV2, and WOX2 (Zhang *et al*., 2017, 2019) (**Supplementary Table S5**). In these clusters, we also found several genes related to suberin accumulation, with some of them even being relevant for periderm development. We identified an AtMyb84 homolog, although not specifically the QsMyb1, two Myb4s, CYP94B1, CYP94B3, and SHR (Almeida *et al*., 2013; Miguel *et al*., 2016; Capote *et al*., 2018; Wang *et al*., 2020; Rojas-Murcia *et al*., 2020; Krishnamurthy *et al*., 2020, 2021; Andersen *et al*., 2021) (**Supplementary Table S5**). Moreover, PER39 (peroxidase), which is involved in proper lignin deposition localization, was also identified (Rojas-Murcia *et al*., 2020) (**Supplementary Table S5**).

Among the most highly transcribed genes related to development and cell wall biogenesis stood out genes related to suberin accumulation (ASFT/FHT, CYP86B1, LTP1.4/LTP2), organ growth (MAT3, XTH, glycosyl hydrolase/endo-1,4 β-D-glucanase, ACAT2, HERK1, RGP), cytokinesis (Ext3), secondary wall of xylem cells formation (glycosyl hydrolase/endo-1,4 β-D-glucanase), xylem differentiation (HB8), ABA signalling pathway (PLDα1) and cell wall integrity (UGD2) (Mishra *et al*., 2006; Drakakaki *et al*., 2006; Cannon *et al*., 2008; Kurasawa *et al*., 2008; Guo *et al*., 2009; Takahashi *et al*., 2009; Compagnon *et al*., 2009; Gou *et al*., 2009; Molina *et al*., 2009; Krupková and Schmülling, 2009; Serra *et al*., 2010; Reboul *et al*., 2011; Jin *et al*., 2012; Chen *et al*., 2016; Deeken *et al*., 2016; Smetana *et al*., 2019) (**Supplementary Table S6**). Moreover, in all these clusters several genes related to suberin (GPAT5, FAR4, KCS2, ABCG2, GELP38, GELP51, GELP96) and lignin accumulation (PER3, PER72) were also recovered (Beisson *et al*., 2007; Franke *et al*., 2009; Lee *et al*., 2009; Domergue *et al*., 2010; Yadav *et al*., 2014; Rojas-Murcia *et al*., 2020; Ursache *et al*., 2021) (**Supplementary Table S6**). Consistent with the upregulation of these genes, several Myb homologs involved in suberin genes induction were found (Myb9, Myb36, Myb84, Myb93, Myb102, and Myc 2) (Kamiya *et al*., 2015; Lashbrooke *et al*., 2016; Legay *et al*., 2016; Capote *et al*., 2018; Wang *et al*., 2020; Wei *et al*., 2020; Wahrenburg *et al*., 2021) (**Supplementary Table S6**). Remarkably in cluster 4, which was enriched in suberin biosynthesis, there were homologs of genes reported to repress suberin accumulation (Myb4, StNAC103/AtNAC058) (Verdaguer *et al*., 2016; Andersen *et al*., 2021). In addition, in this set of clusters, we also found genes previously reported to be related to cambium activity (AIL6, AIL5, WOX4), and phellogen activity (WOX4), as well as to xylem differentiation (LBD18), and phloem differentiation (LBD4) (Yordanov *et al*., 2010; Mudunkothge and Krizek, 2012; Smetana *et al*., 2019; Alonso-Serra *et al*., 2019; Zhang *et al*., 2019; Xiao *et al*., 2020) (**Supplementary Table S6**).

### Cell division, cell expansion and cell differentiation and the bark types

Globally, the gene ontologies enriched in cork-type and rhytidome-type contained upregulated genes that displayed opposite functions referring to cell proliferation, cell expansion, and cell differentiation (**Fig. 5**). These contrasting gene activities align with the phenotype described for cork and rhytidome outer barks, since a major number of larger phellem cells, with high content of suberin, are produced in cork when compared with the rhytidome (Boher et al., 2018). For rhytidome-type barks, we identified upregulated genes related with (i) meristem activity inhibition, (ii) inhibition of cell expansion and (iii) cell differentiation. For example, regarding the most expressed and upregulated genes in rhytidome-type barks, we identified genes that inhibit (i) cell division such as ARF2, BAM3, SVP, PTL and LBD1. ARF2 is a repressor of cell division and flower organ growth (Schruff *et al*., 2006). BAM3 loss of function rescues the root meristem growth in *brx* mutant (Depuydt *et al*., 2013), SVP and PTL inhibit vascular cambium activity (Zhang *et al*., 2019) and PtLBD1 suppresses the vascular cambium cell identity and promotes phloem differentiation (Yordanov *et al*., 2010). In relation to (ii) cell expansion, in rhytidome-type barks we identified upregulated genes that inhibit it. For instance, Feronia reduces cell expansion by binding to RALF (rapid alkalinization factor) and increasing the apoplastic pH (Haruta *et al*., 2014), as well as promotes crosslinking between cell wall pectins by pectin de-esterification (Duan *et al*., 2020). Concerning (iii) cell differentiation, in rhytidome-like barks several positive regulators of triggering cell differentiation over meristematic cell state were upregulated such as LBD1 (mentioned above), BAM3, ARF6, KNAT1/BP, STM, and LBD4. BAM3 was proposed to participate in the differentiation of protophloem (Depuyt *et al*., 2013). ARF6, expressed in all stages of root periderm development in Arabidopsis (Xiao *et al*., 2020), induces vascular patterning and epidermal cell differentiation through negative regulation of class 1 KNOX genes (Tabata *et al*., 2010). About these KNOX genes, we identified KNAT1/BP, that, despite promoting vascular cambial activity (Zhang *et al*., 2019) and increasing the number of periderm cell layers (Xiao *et al*., 2020) in the root, it has opposite role in the hypocotyl by promoting xylem differentiation together with STM, another class I KNOX gene (Liebsch *et al*., 2014), which was also upregulated in rhytidome-type samples. LBD4 was considered a major node in the network of vascular development (Zhang *et al*., 2019) related to phloem recovery defects, possibly acting as a boundary regulator or as an amplifier of divisions on the phloem side of the procambium (Smit *et al*., 2020). Conversely, regarding cork-type barks we identified genes (i) promoting cell division (AIL6, HB8, AIL5, RGP, Ext3, cyclins, and cyclin-dependent kinase) and meristem maintenance (AIL6, glycosyl hydrolase, WOX4, HB8) and, and (ii) some genes involved in cell expansion (XTHs, ACAT2, ERK1, and expansins) and radial growth (LBD4 and LBD18), supporting the superior cell size and cell production of phellem layers in cork oak. Regarding (i) meristem activity, AIL6, together with ANT and AIL7, is required for meristem maintenance, by promoting cell division and repressing cell differentiation in shoot apical meristem (Mudunkothge and Krizek, 2012). A glycosyl hydrolase upregulated is a membrane-bound endo-1,4 β-D-glucanase involved in cellulose synthesis necessary for maintaining meristematic pattern, organ growth in shoot and root and for hormone response (Krupková and Schmülling, 2009) that can regulate cortical microtubule organization (Paredez *et al*., 2008). As concerns to WOX4, it has been shown that it promotes phellogen activity in root periderm (Xiao et al., 2020) and HB8 inhibits cell division and promotes cellular quiescence in the vascular cambium stem-cell organizer, located at xylem side of the vascular cambium but able to maintain xylem and phloem identity at both sides (Smetana *et al*., 2019). These results allow us to speculate that HB8 would induce a similar dynamic organizer within the phellogen stem cell population, which would also accumulate WOX4, as reported for vascular cambium (Smetana *et al*., 2019). Cork-type barks also showed upregulation of genes involved in cell division and/or cell plate formation such as RGP and EXT3 (Drakakaki *et al*., 2006; Cannon *et al*., 2008). As regards (ii) genes inducing cell expansion, we identified various genes in cork-type barks (XTH, ACAT2, HERK1). XTH is able to modify xyloglucans chains, which turnover is required during cell and organ elongation (Kurasawa *et al*., 2008; Yan *et al*., 2019). ACAT2 catalyses the formation of a mevalonate-derived isoprenoids with consequences on the proper growth of vegetative tissues and special effect on cell number in xylem and phloem (Jin *et al*., 2012). HERK1 is a receptor-like kinase (RLKs) shown to be involved in cell expansion by regulating xyloglucan endotransglucosylase/hydrolases and expansins (Guo *et al*., 2009). According to this function, several xyloglucan endotransglucosylase/hydrolase and expansins were also found upregulated in cork-type barks and specifically in the same cluster. Finally, several LOB domain-containing proteins involved in radial growth (Zhang *et al*., 2019) were upregulated in cork-type barks. It was shown that one of them was expressed in secondary phloem (LBD4) and the other expressed in secondary xylem (LBD18) and it was suggested that LBD4 was involved in recruiting cells into the phloem lineage while defining the phloem-procambium boundary (Yordanov *et al*., 2010; Smit *et al*., 2020).

**Fig. 5.**
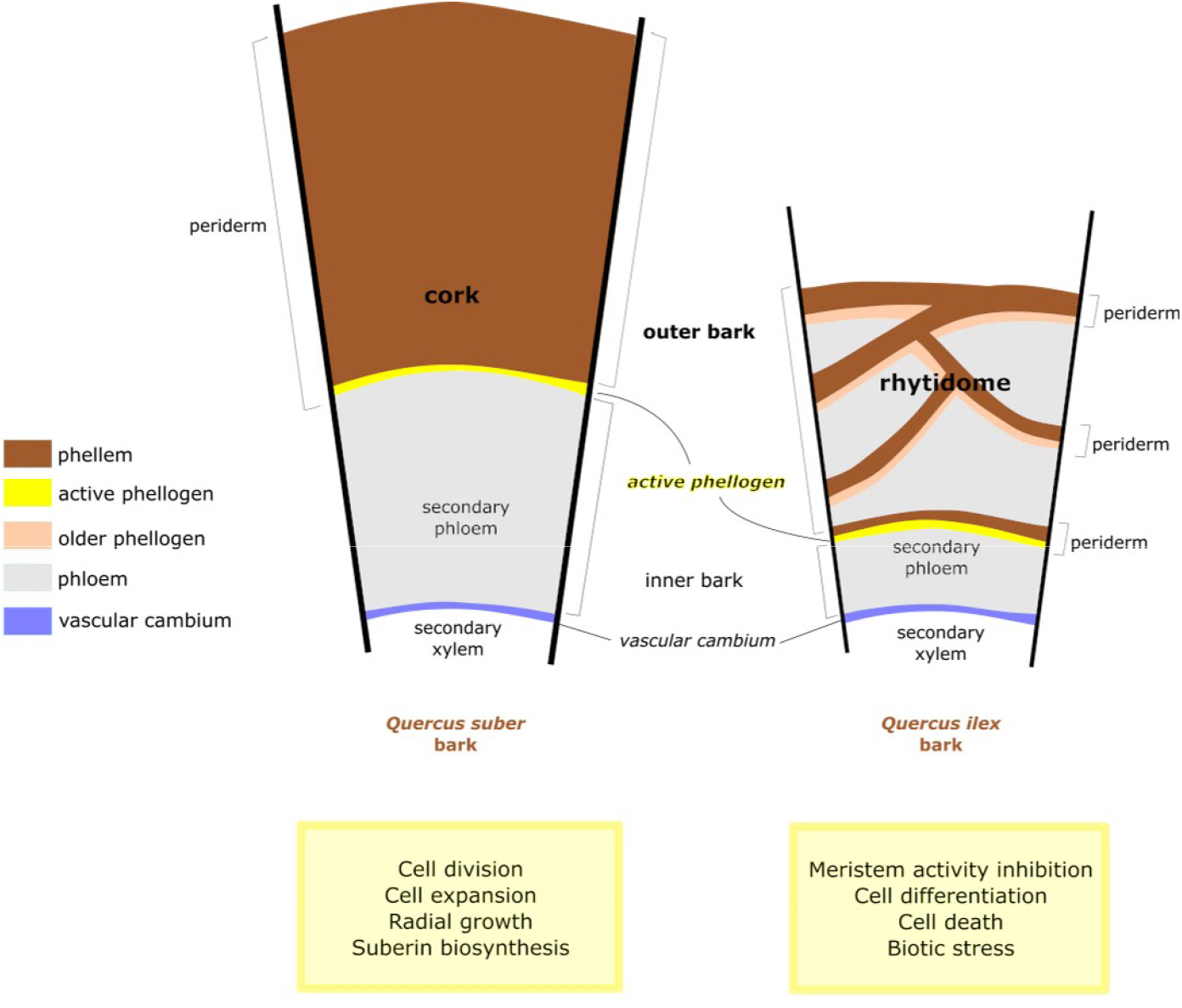
Summary of biological processes occurring during cork and rhytidome formation. This summary is based on upregulated genes and processes in cork-type and rhytidome-type outer barks from *Q. suber, Q. ilex* and their natural hybrids (cork-like and rhytidome-like). The outer tissue portion analysed corresponded to the inner face of the outer bark, which includes the meristematic active cells of phellogen and the alive phellem cells, and for rhytidome-type bark also included alive secondary phloem. Phellogen in *Q. suber* extends concentrically, is reactivated every growing season and forms a persistent periderm during the entire tree life called cork. In *Q. ilex*, the periderm is not persistent and is substituted for new and active phellogens formed inwardly within secondary phloem and yielding a rhytidome outer bark constituted by subsequent periderms with phloem tissue enclosed between them. The phelloderm, derived from each phellogen and located inwardly, has been omitted for simplicity; phelloderm, phellogen and phellem constitute each of the periderms depicted. Sketch inspired from Junikka (1993).

### Cell abscission related processes align with rhytidome-type bark features

One of the most striking differences between rhytidome- and cork-type barks is the shedding of outer layers from rhytidome and the ability to keep one unique persistent periderm within yearly produced phellem cells. It is highly remarkable SOBIR1, upregulated in rhytidome, which was recently suggested to contribute to organ abscission signalling downstream of SERK proteins (Taylor et al., 2019). Organ abscission is a precisely controlled process that gives rise to cell wall loosening and degradation of cell wall components, being pectin-rich middle lamella the major physical mediator of cell adhesion and separation (Daher and Braybrook, 2015). Besides, organ abscission is induced by jasmonic acid, which overlaps with defence processes (Patharkar and Walker, 2018)and lignin deposition also takes place to the abscised region limit to restrict cell wall hydrolyzing enzymes (Lee et al., 2018). It is worth mentioning that cluster 3, induced in rhytidome-type barks and with a peak in rhytidome, in which SOBIR1 is found, is enriched in biotic stimulus, lignin, jasmonic acid, and cell wall biogenesis. Altogether unveil that cell wall related genes identified in this study can be insightful and it is tempting to speculate that cell abscission is an active process leading to rhytidome-type bark.

## CONCLUSION

The main goal of the present study is to provide insight into the molecular mechanisms driving the development of different types of outer bark in woody species, namely the most common rhytidome (characterized by anastomosed thin periderms, encompassing sectors of lignified dead phloem) and the unique, thick phellem typical of *Q. suber* and few other species which present a single long-lived or persistent periderm For this purpose, we have combined chemical, anatomical and transcriptomic approaches in *Q. ilex* (rhytidome), *Q. suber* (thick cork) and hybrid samples. Analysis of the chemical composition of these bark types is consistent with anatomical observations, with *Q. suber* yielding a larger suberin amount, while hybrid samples show different intermediate situations. Inclusion of hybrids has allowed us to highlight 8,336 candidate genes. We confirm that for all outer bark types abiotic stress is a common signal, cork-type barks are enriched in GOs related to lipid metabolism and cell cycle while rhytidome-barks are mostly enriched in GOs related to biotic stress and cell death. Focusing on cell wall biogenesis and development, genes promoting meristem activity and cell expansion are upregulated in cork-type barks, while rhytidome-type barks show higher expression of genes inhibiting cell division and expansion and promoting cell differentiation. Further research is needed in order to disentangle the regulatory pathways of the candidate genes identified in this work, as well as their additive and non-additive effects on bark development.

## Supplementary data

**Table S1:** The amount and quality of RNA, statistics of RNA-Seq data and gene expression profile in outer bark of cork oak, holm oak and the hybrids.

**Table S2**. Primer sequences used for RNA-seq validation by Real Time PCR.

**Table S3**. Chemical composition of outer bark (%) of cork oak (cork, *Quercus suber*), holm oak (rhytidome, *Quercus ilex*) and the *Q. ilex x Q. suber* hybrids. There are five hybrids showing a rhytidome-like bark (FS16 to FS22) and one showing a cork-like bark (FS1).

**Table S4**. GO enrichment analyses for genes included in different clusters (cluster 1 to cluster 8) using the Arabidopsis best homologue. Corrected p-values for False Discovery Rate (FDR) are displayed for each GO term over-represented. The cutoff was set up at FDR ≤ 0.05. Related terms were manually classified into general categories. Term type: P= Biological Process; F= Molecular Function; C= Cellular Component.

**Table S5**. A selection of most expressed genes related to development and found in clusters 1, 2 and 3. Also genes related to periderm development, suberin and lignin accumulation and lateral meristems are included.

**Table S6**. A selection of most expressed genes related to development and found in clusters 4, 5 and 6. Also genes related to periderm development, suberin and lignin accumulation and lateral meristems are included.

**Fig. S1**. Distance map of different outer bark transcriptome profiles. Rhytidome and cork correspond to the outer barks from holm oak and cork oak, and rhytidome-like and cork-like to the outer barks of hybrids. The numbers correspond to the tree identification number.

**Fig. S2**. Gene ontology enrichment for genes found in clusters 1, 2 and 3 with higher expression in rhytidome-type barks. Bars represent the log10 p-value for each GO term. The GO terms were manually compared and those showing the analogous description and same set of genes were grouped, the log10 p-value corresponds to the broader GO term (including the maximum number of genes). The terms are biological process (yellow), molecular function (green), and cell component (blue).

**Fig. S3**. Gene ontology enrichment for genes found in clusters 4, 5, 6, 8 with higher expression in cork-type barks. Bars represent the log10 p-value for each GO term. The GO terms were manually compared and those showing the analogous description and same set of genes were grouped, the log10 p-value corresponds to the broader GO term. The terms are biological process (yellow), molecular function (green), and cell component (blue).

**Fig. S4**. Gene ontology enrichment for genes found in cluster 7 with particular higher expression in cork-like and rhytidome barks. Bars represent the log10 p-value for each GO term. The GO terms were manually compared and those showing the analogous description and same set of genes were grouped, the log10 p-value corresponds to the broader GO term. The terms are biological process (yellow), molecular function (green), and cell component (blue).

## Acknowledgements

The authors are very grateful to Sandra Fernández-Piñán, Jennifer López, Francisco Martínez Moreno and Antonio Rodríguez for harvesting outer barks.

## Author contribution

ULH, MS, AS, OS and MF conceived and designed the experiment; ULH and AS performed anatomical observations; IA, MS, AP, OS and MF performed the RNA extraction; IA and ULH performed the bioinformatics analyses; MR and PJ performed the chemical analyses of outer barks; AP performed qPCR; IA and MF interpreted the data, which were discussed with OS, AS and ULH. IA and MF wrote the manuscript and IA, MF, OS, AP, ULH and AS made the figures. All authors revised the final manuscript form.

## Data availability statement

All data supporting the findings of this study are available within the paper, within its supplementary materials published online and in the Gene Expression Omnibus repository from NCBI under accession code GSE227020.

